# Human coronavirus OC43 infection remodels Connexin 43 mediated gap junction intercellular communication *in vitro*

**DOI:** 10.1101/2023.11.20.567927

**Authors:** Souvik Karmakar, Jayasri Das Sarma

## Abstract

β-coronaviruses cause acute infection in the upper respiratory tract, resulting in various symptoms and clinical manifestations. OC43 is a human β-coronavirus that induces mild clinical symptoms and can be safely studied in the BSL2 laboratory. Due to its low risk, OC43 can be a valuable and accessible model for understanding β-coronavirus pathogenesis. One potential target for limiting virus infectivity could be gap junction-mediated communication. This study aims to unveil the status of cell-to-cell communications through gap junctions in human β-coronavirus infection. Infection with OC43 leads to reduced expression of Cx43 in A549, a lung epithelial carcinoma cell line. Infection with this virus also showed a significant ER and oxidative stress increase. Internal localization of Cx43 is observed post OC43 infection in the ERGIC region, which impairs the gap junction communication between two adjacent cells, confirmed by Lucifer yellow dye transfer assay. It also affects hemichannel formation, as depicted by the EtBr uptake assay. Altogether, these results suggest that several physiological changes accompany OC43 infection in A549 cells and can be considered an appropriate model system for understanding the differences in gap junction communication post-viral infections. This model system can provide valuable insights for developing therapies against human β-coronavirus infections.

**Importance:** The enduring impact of the recent SARS-CoV-2 pandemic underscores the importance of studying human β-coronaviruses, advancing our preparedness for future coronavirus infections. Due to SARS-CoV-2 being highly infectious, another human β-coronavirus OC43 can be considered as an experimental model. One of the crucial pathways that can be considered is gap junction communication, as it is vital for cellular homeostasis. Our study seeks to understand the change in Cx43-mediated cell-to-cell communication during human β-coronavirus OC43 infection. *In vitro* studies showed the downregulation of the gap junction protein Cx43 and the upregulation of endoplasmic reticulum and oxidative stress markers post-OC43 infection. Furthermore, OC43 infection causes impairment of functional hemichannel and gap junction formation. Overall, this current study infers that OC43 infection reshapes intercellular communication, suggesting that this pathway may be a promising target for designing highly effective therapeutics against human coronaviruses by regulating Cx43 expression.

## Introduction

Gap junction intercellular communication (GJIC) maintains cellular homeostasis in multicellular organisms by allowing cells to coordinate and respond to signals. Gap junctions are specialized channels that exchange ions, signaling molecules, and small molecules (especially less than 1 kDa) between adjacent cells [1]. This connection aids in rapid and synchronized cellular responses critical for various physiological processes. Gap junctions are formed with protein subunits called connexons, comprising a hexameric assembly of connexins. When present on the cell surface, these connexons develop hemichannels that make these cells susceptible to external stimuli. The coupling of the connexon from one cell with the connexon of another cell generates functional gap junctions [2]. These gap junctions in their open conformations provide a passage between adjacently present cells that facilitates the passive diffusion of essential molecules to maintain cellular homeostasis [3].

The expression of gap junction proteins, particularly connexin, has been reported to be modulated following virus infections [4]–[6]. Connexin 43 (Cx43), the most ubiquitously expressed and widely studied among all connexins, has been extensively studied in response to murine β-coronavirus infections [7]–[10]. The regulation in Cx43 following virus infection is attributed to either viral burden on the endoplasmic reticulum (ER) causing cellular stress or hijacking microtubule-mediated transport for trafficking of viral proteins within the cells. Virus burdens the endoplasmic reticulum of the host cell by translating a large number of viral proteins [11]. Virus-induced ER stress can alter host cellular processes, potentially causing cellular damage. ER stress can be detected by different host cell markers, including heat shock factor (HSF1), heat shock protein 70 (HSP70), and endoplasmic reticulum protein 29 (ERp29). Under ER stress, HSF1 regulates the production of HSP70, a chaperone, by binding to the heat shock elements (HSE)[12]. HSP70 stabilizes unfolded proteins through selective binding, thus preventing aggregate formation while parallelly facilitating refolding of misfolded proteins[13], [14]. ERp29 is a major chaperone protein that helps in the trafficking of membrane proteins to the cell surface, including Cx43 [15]. In addition to its role in protein trafficking, overexpression of ERp29 induced by ER stress significantly contributes towards overall unfolded protein stress. More specifically, ERp29-mediated activation of protein-degrading machinery aids in the clearance of denatured proteins and prevents protein aggregation[16]. Recent evidence has demonstrated that viral infections are coupled with a rise in the production of reactive oxygen species (ROS), which the host cannot combat. Resultant oxidative stress generated in the host cell can be detected by regulation of Parkinson disease protein 7, DJ-1. DJ-1 has roles in transcriptional regulation and antioxidative stress reaction, which helps recover from oxidative stress [17], [18].

This study employs the HCoV-OC43 virus to elucidate the effect of viral infection in modulating gap junction-mediated cell-to-cell communication in a human lung epithelial cell line, A549. OC43 is a human β-coronavirus that causes mild symptoms in humans and hence can be a suitable model for understanding the pathogenesis caused by the infections of human coronaviruses. OC43 shares many similarities with respect to genome size, transmission mode, common symptoms, and pathogenesis with other human β-coronaviruses, including SARS-CoV-2. Studies on human coronaviruses are needed because of the lack of available human data and the reproducibility of pathogenesis in other virus models. Hence, studying β-coronaviruses in human models is essential, and OC43 can be an excellent option. Here, we used a human lung epithelial cell line, A549, to understand the changes in functional gap junction and hemichannel formation post-OC43 infection. We observed that post-infection Cx43 is reduced at the protein level, accompanied by a rise in ER and oxidative stress markers. Cx43 is retained in the perinuclear region of infected cells, confirmed by immunofluorescence and Triton X-100 solubilization assay. With the decrease in Cx43, the cells could not form functional gap junctions and hemichannels, confirmed by the Lucifer yellow dye transfer assay and the Ethidium Bromide uptake assay. These data suggest that targeting Cx43 to modulate its expression can be a valuable therapeutic target to reduce β-coronavirus infections.

## Materials and Methods

### Reagents

Bicinchoninic acid (BCA) protein assay kits and Super Signal West Pico Chemiluminescent Substrate were procured from Thermo Fisher Scientific. Protease and phosphatase inhibitor cocktails were from Sigma-Aldrich. Cell culture reagents, including Dulbecco’s modified Eagle’s medium (DMEM), fetal bovine serum (FBS), 0.25% trypsin, and Penicillin/streptomycin, were obtained from Gibco. Primary antibodies used in the study are listed in Table 1, and the secondary antibodies used are listed in Table 2. The remaining chemicals were from Merck (Deu, Germany), SRL (MB, India), Sigma-Aldrich (MO, USA), or Thermo Fisher Scientific (MA, USA).

**Table 1.**
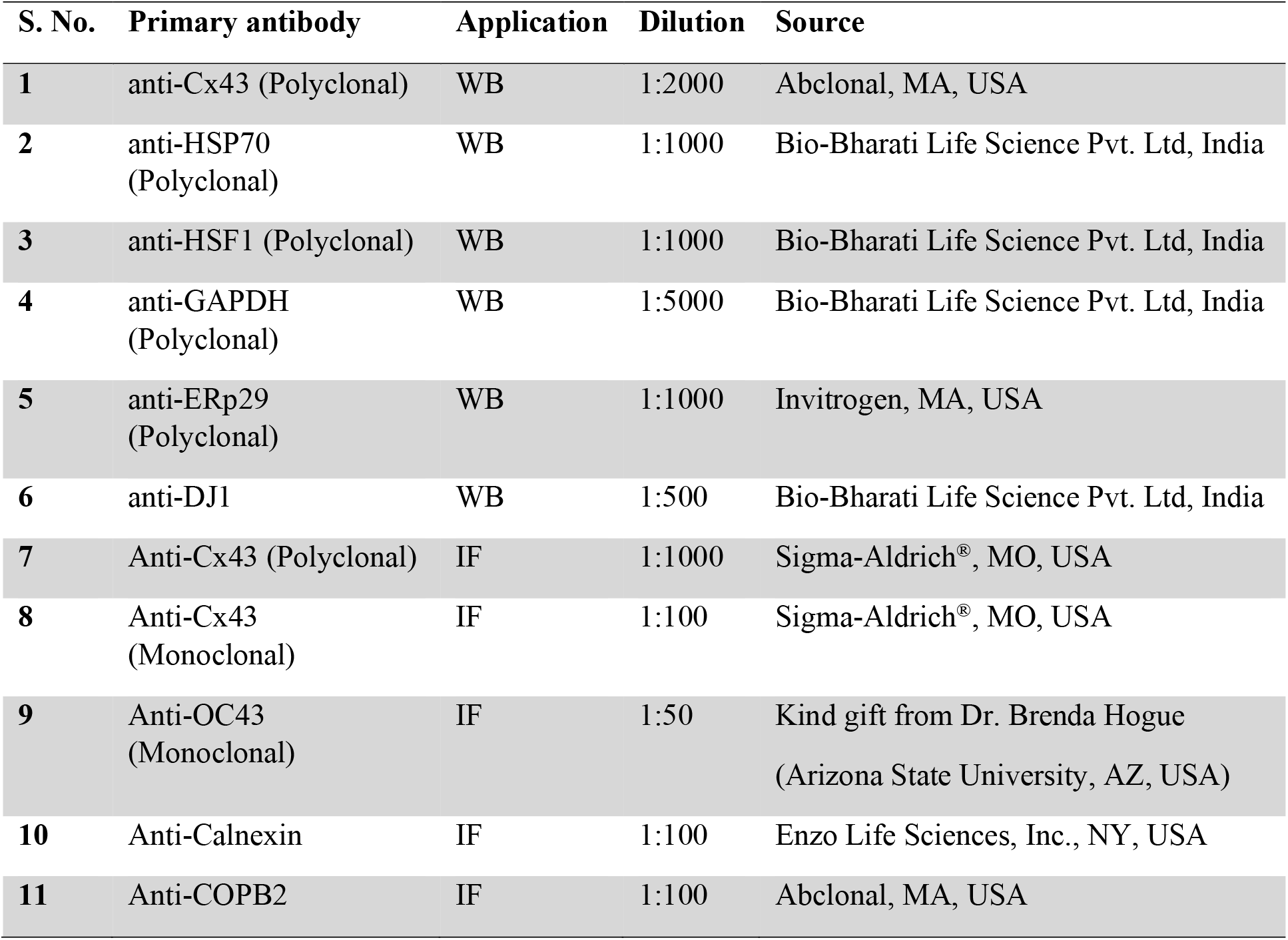
Primary antibodies used in this study.

| S. No. | Primary antibody | Application | Dilution | Source |
| --- | --- | --- | --- | --- |
| 1 | anti-Cx43 (Polyclonal) | WB | 1:2000 | Abclonal, MA, USA |
| 2 | anti-HSP70 (Polyclonal) | WB | 1:1000 | Bio-Bharati Life Science Pvt. Ltd, India |
| 3 | anti-HSF1 (Polyclonal) | WB | 1:1000 | Bio-Bharati Life Science Pvt. Ltd, India |
| 4 | anti-GAPDH (Polyclonal) | WB | 1:5000 | Bio-Bharati Life Science Pvt. Ltd, India |
| 5 | anti-ERp29 (Polyclonal) | WB | 1:1000 | Invitrogen, MA, USA |
| 6 | anti-DJ1 | WB | 1:500 | Bio-Bharati Life Science Pvt. Ltd, India |
| 7 | Anti-Cx43 (Polyclonal) | IF | 1:1000 | Sigma-Aldrich®, MO, USA |
| 8 | Anti-Cx43 (Monoclonal) | IF | 1:100 | Sigma-Aldrich®, MO, USA |
| 9 | Anti-OC43 (Monoclonal) | IF | 1:50 | Kind gift from Dr. Brenda Hogue (Arizona State University, AZ, USA) |
| 10 | Anti-Calnexin | IF | 1:100 | Enzo Life Sciences, Inc., NY, USA |
| 11 | Anti-COPB2 | IF | 1:100 | Abclonal, MA, USA |

**Table 2.** Secondary antibodies used in this study.

| S. No. | Secondary Antibody | Application | Dilution | Source |
| --- | --- | --- | --- | --- |
| 1 | Alexa 488 Goat Anti-Rabbit | IF | 1:1000 | Invitrogen, MA, USA |
| 2 | Alexa 488 Goat Anti-Mouse | IF | 1:1000 | Invitrogen, MA, USA |
| 3 | Alexa 568 Donkey Anti-Rabbit | IF | 1:1000 | Invitrogen, MA, USA |
| 4 | Alexa 568 Donkey Anti-Mouse | IF | 1:1000 | Invitrogen, MA, USA |
| 5 | HRP conjugated Rabbit secondary antibody | WB | 1:10000 | Invitrogen, MA, USA |

### Cells and viruses

Human epithelial lung carcinoma cell line A549 was obtained from Dr. Michael Koval (Emory University School of Medicine, Atlanta, Georgia, USA) and African green monkey kidney epithelial cells, Vero cells, a kind gift from Dr. Sourish Ghosh (Indian Institute of Chemical Biology, Kolkata, West Bengal, India). The cell lines were maintained by DMEM medium supplemented with 10% heat-inactivated FBS and 1% penicillin and streptomycin. The cells were maintained at 37⁰C in a humidified incubator with 5% CO2. Human coronavirus OC43 (HCoV-OC43) was obtained through BEI Resources, NIAID, NIH (catalog number: NR-56241). The virus was propagated in Vero cells for further use.

### Virus propagation and virus infection

For virus propagation, a 60mm dish of around 80% confluent Vero cells was inoculated with OC43 at a multiplicity of infection (MOI) of 1 in 500 µl of infection media (DMEM supplemented with 2% FBS and 1% penicillin and streptomycin). The virus was allowed to adsorb onto the cells for 2 hours with intermediate shaking after 15-minute intervals at 33⁰C in a humidified incubator with 5% CO2. The infection media was removed after 2 hours, and 2.5ml of infection media was added. The plate was incubated for four days at 33⁰C with 5% CO2. Four days were chosen for virus incubation based on the cytopathic effects exhibited by the virus on Vero cells, which is discussed later. The media was collected from the dish and centrifuged at 500g for 5 minutes to clarify the viral particles. The supernatant obtained was aliquoted and kept at -80⁰C for further use. For virus infection, 80% confluent A549 cells were inoculated with OC43 at MOI 0.5 and 1 in infection media. The virus inoculum was adsorbed for 2 hours at 37⁰C, 5% CO2, with shaking every 15 minutes. The inoculum was removed after 2 hours, and fresh infection media was added. The cells were incubated at 37⁰C with 5% CO2.

### Viral titer determination by TCID50 assay

Vero cells were plated in a 96-well plate and incubated at 37⁰C with 5% CO2 till the cells were 70-80% confluent. The media was removed from the wells, and a PBS wash was given. The PBS was removed, and the cells were inoculated with 10-fold serial dilutions of OC43 stock virus. 100 µl of the dilutions were given to each well, and the plate was incubated for six days at 37⁰C with 5% CO2. After six days, the plate was removed, and the media was discarded. At room temperature, the cells were fixed in 4% paraformaldehyde (4% PFA) for 15 minutes. The cells were permeabilized with 0.2% Triton X-100 in PBS after fixation. 50 µl of blocking solution was given using 2.5% heat-inactivated goat serum in PBS with 0.2% Triton X-100. The incubation was carried out for 15 minutes at room temperature. 25 µl of primary antibody diluted in blocking solution was added to the wells and was incubated for 2 hours at room temperature. Similarly, after washing, 25 µl of secondary antibody (Alexa Fluor 488) diluted in blocking solution was given to the wells and incubated for 1 hour 15 minutes at room temperature. The wells were washed, and images were taken using a 10X objective in a Nikon Eclipse Ts2-FL microscope (Tokyo, Japan) and device camera Nikon DS-Fi3 (Tokyo, Japan). The Reed-Muench method was used for the calculation of TCID50 [19]. To convert the TCID50/mL values to PFU/mL, TCID50/mL values were multiplied by 0.7 [20].

### Immunofluorescence

A549 cells were grown on etched coverslips, and virus infection was performed. After infection, cells were washed with wash buffer (PBS with calcium and magnesium). 4% PFA was used for cell fixation for 10 minutes at room temperature. To permeabilize the cells, PBS, along with 0.2% Triton X-100, was used. Following permeabilization, cells were blocked using a blocking solution made of PBS with 0.2% Triton X-100. The primary antibody was diluted in the blocking solution, and 30 µl drops were applied on a surface where the coverslips were inverted. The setup was placed in a humidified chamber overnight at 4⁰C. After washing, the coverslips were similarly placed on drops of secondary antibody and incubated for 1 hour 15 minutes at room temperature. Cells were washed and mounted in Mowiol containing DAPI to stain the nucleus. The slides were imaged in an Olympus IX-81 microscope and a Zeiss Confocal Laser Scanning microscope (LSM710). Images were processed with ImageJ (NIH) and Zen 3.7 (Carl Zeiss AG) software.

### Protein isolation and immunoblotting

Cultured cells were subjected to virus infections and incubated for three days. After three days, the media was discarded and rinsed twice with ice-cold PBS. RIPA buffer was used for cell lysis which comprised 50 mM Tris, 150 mM NaCl, 0.1% sodium dodecyl sulfate (SDS), 0.5% sodium Deoxycholate (C24H40O4), 1% Triton X-100, EDTA-free complete protease and phosphatase inhibitors (1 mM sodium orthovanadate, 10 mM sodium fluoride, and 10 mM sodium pyrophosphate decahydrate). The cell lysis process was conducted on ice for 1 hour and 30 minutes with intermittent vortexing every 15 minutes. Centrifugation at 13,200 rpm for 20 minutes at 4⁰C was performed using an Eppendorf 5415 R centrifuge to clarify the cell lysate. The pellet was discarded, and the supernatant was collected as it contained the cellular proteins and was estimated for total protein concentration. It was determined using a Pierce® BCA protein assay kit (Thermo Scientific, Rockford, IL, USA) following the manufacturer’s instructions. An equal amount of protein (10 µg) was subsequently separated by 12% SDS-PAGE and then transferred onto a polyvinylidene difluoride (PVDF) membrane (Millipore, Bedford, MA). The membrane was blocked with a 5% skimmed milk solution prepared in Tris-buffered saline containing 0.1% Tween-20 (TBS-T) for 1 hour at room temperature. The membrane was then incubated overnight at 4⁰C with primary antibody solutions diluted in the blocking solution. After the primary antibody incubation, the membrane was washed thrice with TBS-T for 10 minutes each. Subsequently, it was incubated at RT for 1 hour with an appropriate secondary antibody conjugated with horseradish peroxidase (HRP) prepared in a blocking solution. After washing with TBS-T, immunoreactive bands were visualized using the Super Signal West Pico Chemiluminescent Substrate (Thermo Scientific) following the manufacturer’s instructions. Images of the blots were captured with the help of GENESys software (Genesys, Cambridge, UK) with the Syngene G: Box ChemiDoc system. The membranes were re-probed with anti-GAPDH antibody to ensure equal protein loading. Densitometric analysis of the obtained bands was performed using ImageJ.

### Triton X-solubilization assay

Cells were collected by scraping in PBS containing protease and phosphatase inhibitors. The cell suspension obtained was passed through a Dounce homogenizer 100 times. The cell homogenate was given a short centrifuge for 5 minutes at 500g using an Eppendorf 5415 R centrifuge, maintaining a temperature of 4⁰C. The supernatant was collected in ultracentrifuge tubes and centrifuged at 100,000g for 30 minutes, maintaining the temperature at 4⁰C using a ThermoFisher Sorvall WX 100+ ultracentrifuge with ThermoFisher TFT-80.2 fixed angle rotor. The membrane-enriched pellet obtained was resuspended in ice-cold PBS containing 1% Triton X-100 and was incubated for 30 minutes, strictly maintaining 4⁰C. After incubation, the samples were centrifuged at 100,000g for 30 minutes at 4⁰C. The supernatant obtained corresponds to the Triton X-100 soluble fraction, and the pellet obtained was considered insoluble. The equal volumes of soluble and insoluble fractions, diluted in Laemmli sample buffer, were resolved by SDS-PAGE followed by Cx43 immunoblotting.

### Lucifer yellow dye transfer assay

Cultured, mock, and infected cells grown in 12-well plates were washed with PBS. A uniform release of 10 µl of Lucifer yellow dye (4 mg/mL concentration) was performed while generating a scratch in the cell monolayer to perform the scrape loading assay [7]. The plate was incubated at room temperature for one minute, followed by a PBS wash. Subsequently, PBS was added to the wells, and the plate was incubated at 37⁰C for 10 minutes to facilitate the diffusion of the loaded dye to the neighboring cells. After incubation, cells were fixed using 4% PFA for 10 minutes at room temperature. Images were captured using an Olympus IX-81 microscope operating at 10X magnification with 488 nm excitation. The distance traveled by the dye was measured with the help of ImageJ (NIH, USA) software [21].

### Ethidium Bromide (EtBr) dye uptake assay

Cells cultured on etched coverslips in a 24-well plate were washed twice with Locke’s solution, comprising 154 mM NaCl, 5.4 mM KCl, 2.3 mM CaCl2, and 5 mM HEPES buffer at a pH of 7.4. Following the washes, a 5 μm Ethidium Bromide (EtBr) solution prepared in Locke’s solution was added to the wells [22]. The plate was incubated for 10 minutes at 37⁰C. The cells were fixed with 4% PFA for 10 minutes at room temperature. Subsequently, a wash was given with Locke’s solution, and the coverslips containing cells were mounted in a Mowiol mounting medium that did not contain DAPI. Images were captured using an Olympus IX-81 microscope operating at 40X magnification. The intensity of EtBr uptake by cells was measured with the help of ImageJ (NIH, USA) software [21].

### Data and statistical analyses

The data from all experiments are presented in the form of mean values accompanied by their standard error of the mean (mean ± SEM). Statistical analysis was performed using a two-tailed student’s t-test with Welch’s correction. Statistical significance was attributed to results with a P value of <0.05. GraphPad Prism 8 software (GraphPad Software, Inc.) was employed for all statistical analyses.

## Results

### Viral growth kinetics in Vero and A549 cells

To determine the optimal virus yield during virus propagation, Vero cells were infected with OC43 at MOI 1 and observed every day till 4 days post-infection. Vero cells developed cytopathic effects (CPE) in the form of cell vacuolization and rounding from day 2 p.i. and continued till day 4 p.i. (Fig. 1B-D). The cells started dying at day 4 p.i. Hence, this time point was chosen as the optimum time for virus propagation.

**Figure 1.**
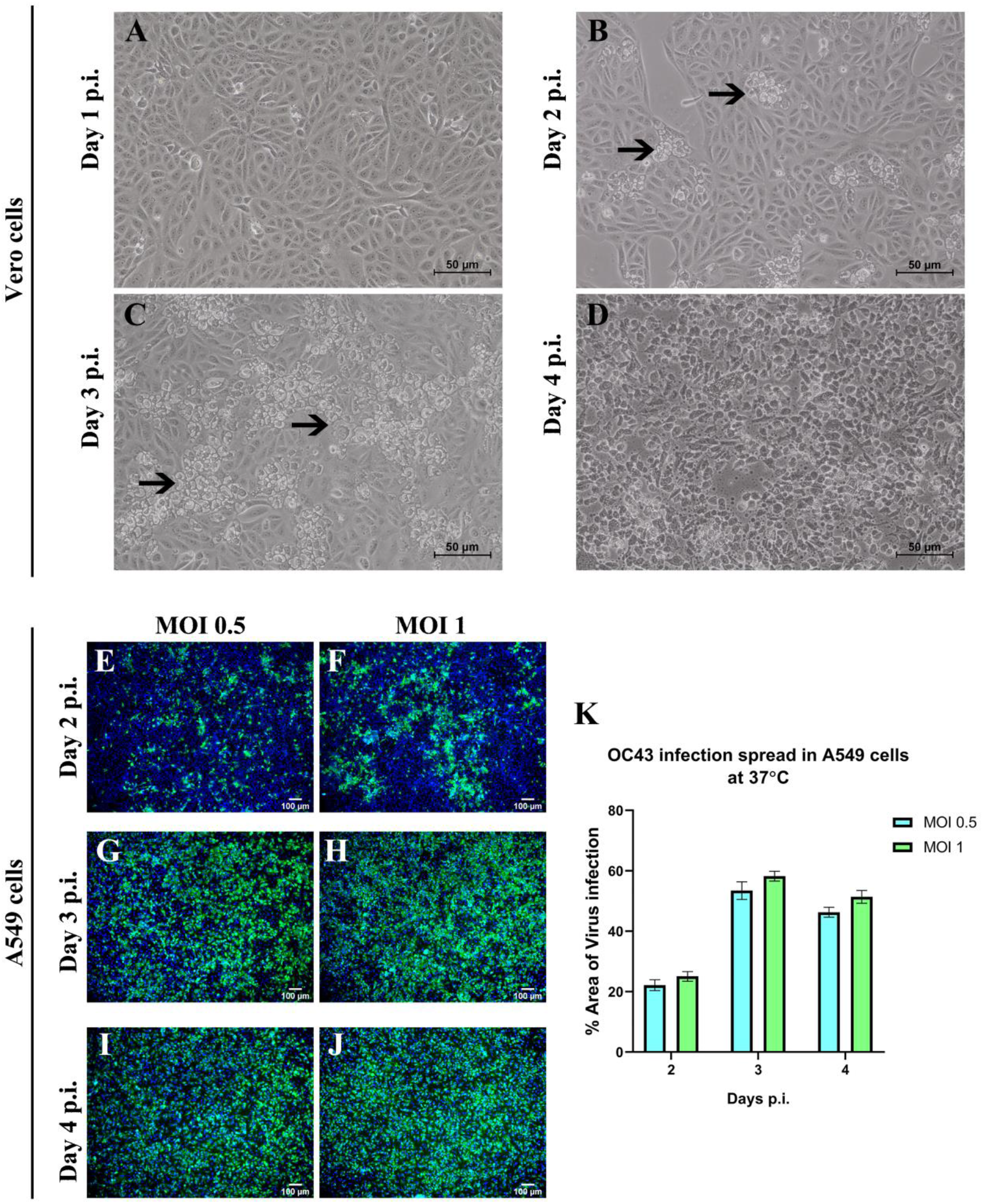
Viral infection in Vero and A549 cells on different days p.i. OC43 infection in Vero cells results in cell vacuolation, depicted with black arrows at different days post-infection (A-D). Infection with OC43 in A549 cells was standardized based on the immunofluorescence staining of virus spread. A549 cells were infected with MOI 0.5 and 1 of OC43, fixed at different time points, immunostained, and imaged in an IX-81 microscope (E-J). The infection spread was quantified using Image J and plotted using the absolute percentage area of virus infection (K). Data expressed as mean± SEM. The scale bar of the images (A-D) depicts 50 microns, and the images (E-J) depict 100 microns.

For investigating the viral infection and spread in A549 cells, the cells were infected with OC43 at MOIs 0.5 and 1, and immunofluorescence with anti-OC43 antibody was performed at days 2, 3, and 4 p.i. (Fig. 1E-J). Day 3 p.i. showed maximum virus infection spread, and on day 4p.i., some of the cells started to die; hence, the infection spread is comparatively lower than on day 3 p.i. as quantified using ImageJ (Fig. 1K). A similar extent of infection was observed in the case of MOI 0.5 and 1 on day 3 post-infection and were chosen for further experiments.

### Reduced Cx43 protein expression post OC43 virus infection

Immunofluorescence imaging and immunoblotting were performed to determine the effect of OC43 infection on gap junction protein Cx43 expression in A549 cells. For immunofluorescence, A549 cells grown on etched coverslips were infected with OC43 at different MOIs of 0.1 and 0.5. Immunostaining was performed at day 3p.i. using rabbit anti-Cx43 and mouse anti-OC43 antibodies. The images were captured in a Zeiss Confocal Laser Scanning Microscope (LSM710) (Fig. 2A-L). It is observed that non-infected cells express punctate staining of Cx43 throughout, depicted with white arrows. The infected cells, depicted with yellow arrows, show reduced punctate staining, suggesting an overall downregulation in Cx43 expression. To confirm the Cx43 expression level alteration, immunoblotting was performed with rabbit anti-Cx43 antibody, and the membrane was stripped and re-probed with rabbit anti-GAPDH antibody to ensure equal loading (Fig. 2M). With the help of densitometric analysis used in ImageJ, it is observed that OC43 infection affects the expression of Cx43 (Fig. 2N). This suggests that upon OC43 infection in A549 cells, the gap junction communication protein Cx43 is reduced.

**Figure 2.**
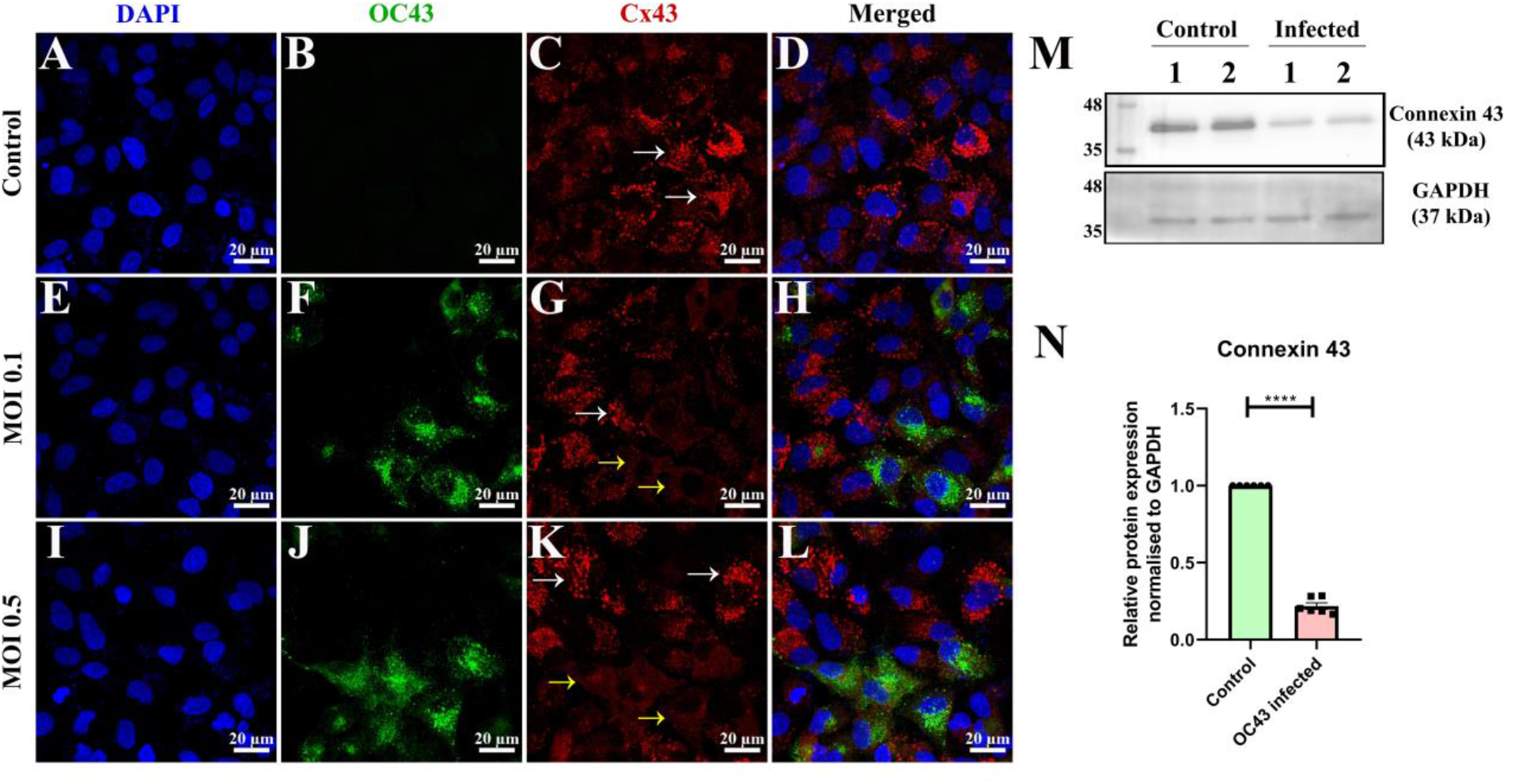
Connexin 43 expression in control and HCoV-OC43 infected A549 cells. A549 cells were immunostained with anti-OC43 antibody and anti-Cx43 antibody, counterstained with DAPI. Images correspond to different sets, such as control (A-D), OC43 infected MOI 0.1 (E-H), and OC43 infected MOI 0.5 (I-L). The non-infected cells showing punctate staining of Cx43 are marked with white arrows (C, G, and K), and the OC43 infected cells showing reduced Cx43 punctate staining are marked with yellow arrows (G and K). D, H, and L represent merged images of the different channels. A western blot determined the overall Connexin 43 expression (M). Densitometric analysis using Image J of the bands obtained was plotted (N). Data expressed as mean± SEM of two individual replicates. The significant differences are indicated by *P<0.05, **P<0.01, ***P<0.001, ****P<0.0001. The scale bar for all the images is 20 microns.

### HCoV-OC43 infection in A549 cells increases ER and oxidative stress

A549 cells were infected with MOI 1 and incubated for 3 days p.i. Total protein was isolated and estimated using BCA assay. 10 µg of protein was loaded in each well for control and OC43-infected samples. We checked the ER stress and oxidative stress in control and infected samples. HSF1, HSP70, and ERp29 were chosen as ER stress markers, and DJ-1 was selected as the oxidative stress marker. As observed by the upregulation of ER stress markers, infection with HCoV-OC43 in A549 cells increased ER stress (Fig. 3A-D). The oxidative stress marker, DJ-1, shows a trend in upregulation, though statistically non-significant (Fig. 3A, E). These results suggest that upon OC43 infection, there is an overall increase in cellular stress.

**Figure 3.**
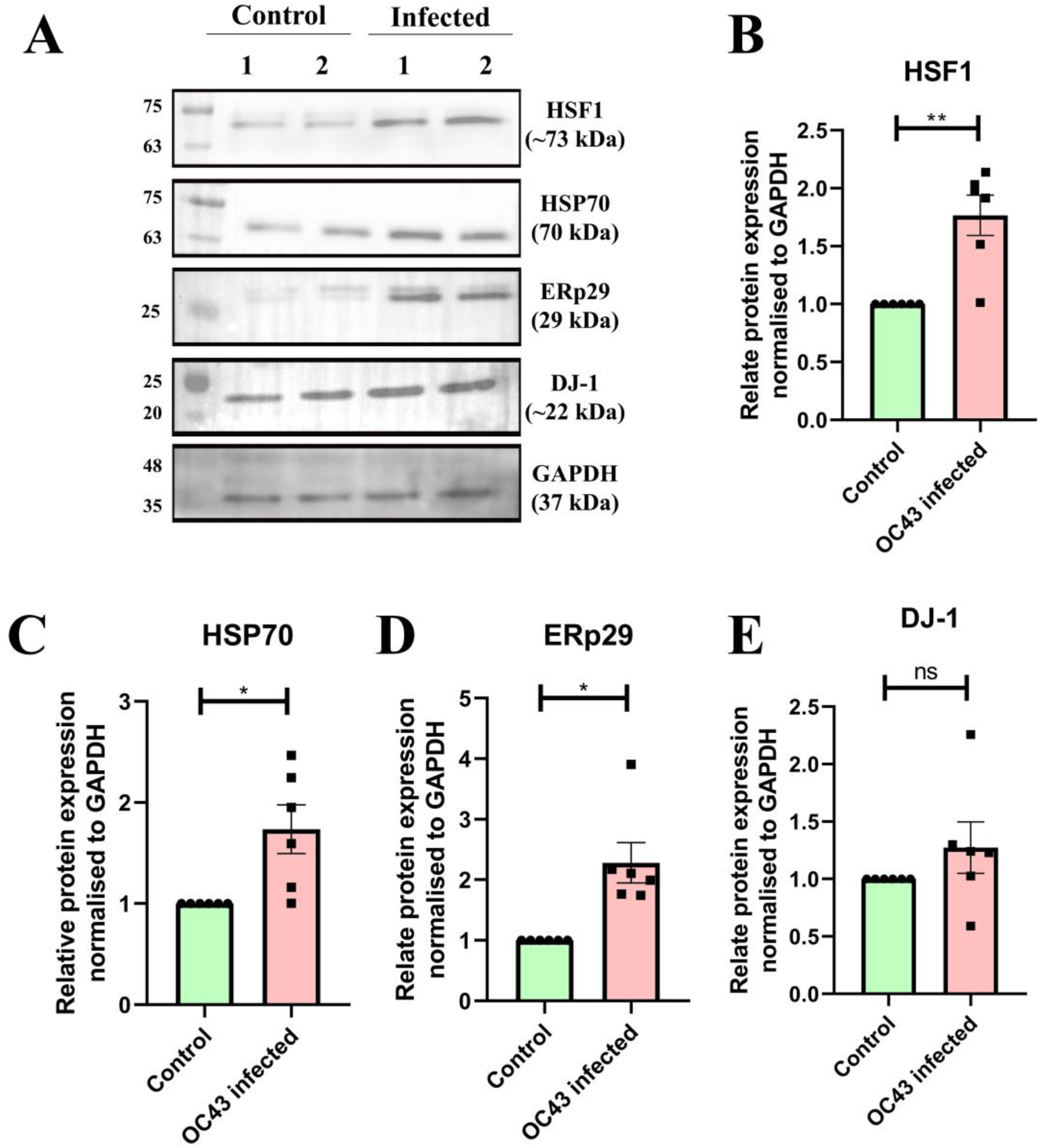
HCoV-OC43 induces ER stress and oxidative stress in A549 cells. Representative immunoblots show the upregulation of different stress markers in infected samples compared to control (A). Scatterplots representing the densitometric analysis of the different stress markers normalized to the loading control GAPDH (B-E). (B) represents significant upregulation of HSF1, (C) represents upregulation of HSP70, (D) represents significant upregulation of ERp29, and (E) represents a trend in upregulation of DJ-1. Data expressed as mean± SEM of two individual replicates. The significant differences are indicated by *P<0.05, **P<0.01, ***P<0.001, ****P<0.0001.

### Internal localization of Cx43 upon HCoV-OC43 infection

We determined the localization of Cx43 in mock and infected cells. A549 cells, grown on etched coverslips, were infected with OC43 and incubated for 3 days. Immunofluorescence staining was performed using rabbit anti-Cx43 and mouse anti-OC43 antibodies. Images were taken in an IX-81 microscope (Fig. 4A-H). An overall Cx43 punctate staining was observed in the control cells, depicted by white arrows (Fig. 4D). The yellow arrow depicts the infected cells with reduced punctate staining and perinuclear localization (Fig. 4H). The reduction of Cx43 in gap junctional plaques upon virus infection was confirmed by the Triton X-100 solubilization assay. The densitometric analysis of infected samples showed a significant decrease in the ratio of triton X-100 insoluble to soluble levels of Cx43 when compared to control samples (Fig. 4I, J). This further indicates increased Cx43 gap junction assembly at the cell surface in mock-infected samples compared to infected samples.

**Figure 4.**
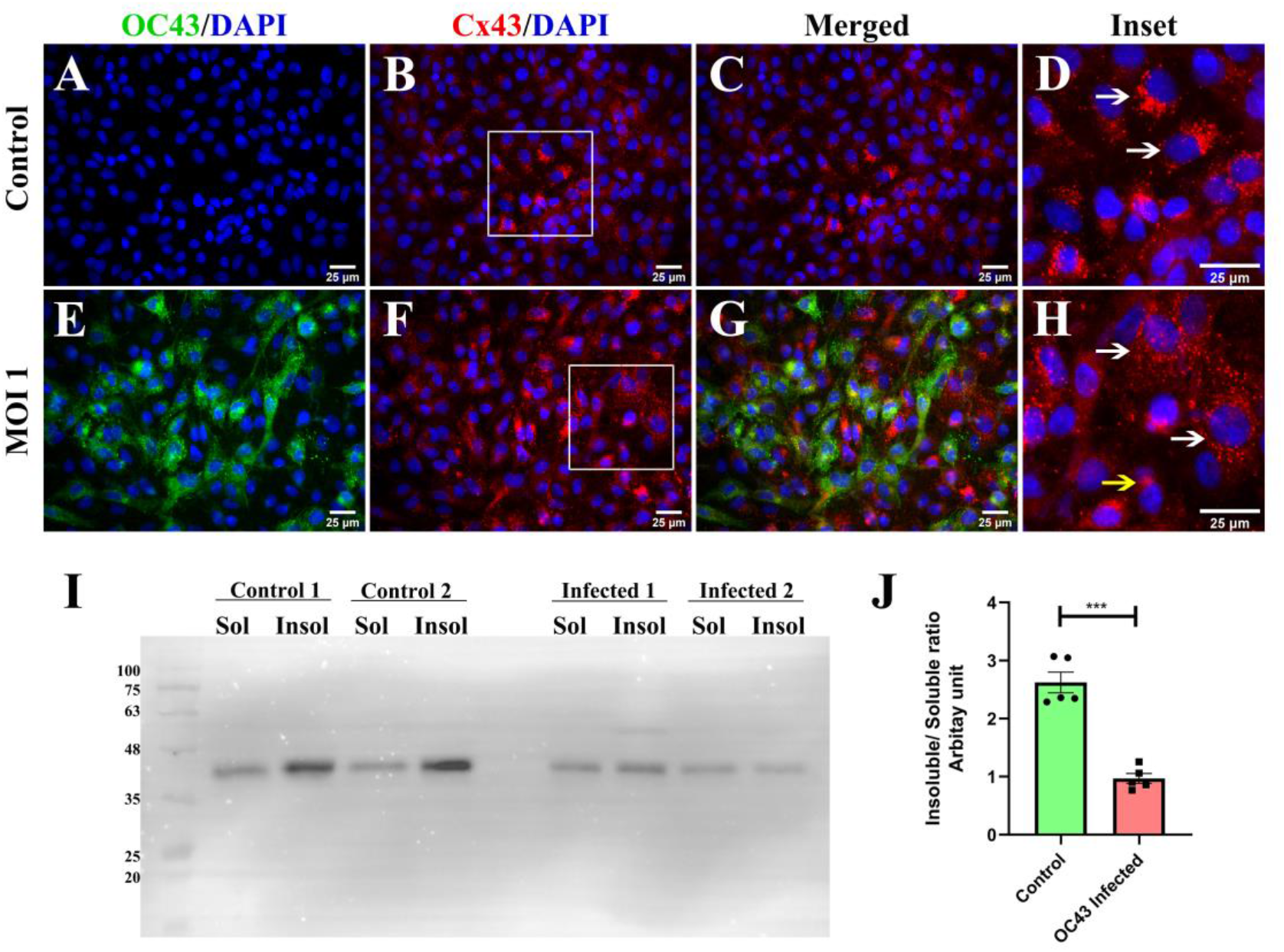
HCoV-OC43 infection in A549 cells impairs Cx43 trafficking to the cell surface. A549 cells control (A-C) and infected (E-G) were immunostained with anti-OC43 and anti-Cx43 and counterstained with DAPI. The control cells show punctate staining of Cx43 throughout the cell, depicted by white arrows (D). In contrast, infected cells show reduced Cx43 punctate staining and internal localization of Cx43 in the perinuclear region depicted by the yellow arrow (H). To confirm the internal localization of Cx43 upon OC43 infection, a Triton X-100 solubilization assay was performed (I), and the densitometric analysis of the bands obtained revealed that the Insol/sol fraction of control is greater than the infected samples (J). The insets are digitally zoomed. The scale bar for all the images is 25 microns. Data expressed as mean± SEM of three individual replicates. The significant differences are indicated by *P<0.05, **P<0.01, ***P<0.001, ****P<0.0001.

To identify the subcellular localization of Cx43 upon OC43 infection, we performed immunostaining of an ER resident protein, Calnexin, using rabbit anti-calnexin antibody and ERGIC marker, COPB2, using rabbit anti-COPB2 antibody. Confocal images showed that infected cells have perinuclear connexin staining (Fig. 5B, F, J, and N). The control and non-infected cells are marked with white arrows (Fig. 5D, H, L, and P), and the infected cells are marked with yellow arrows (Fig. 5H and P). Visual analysis of the microscopic images reveals that the perinuclear localization of Cx43 in infected cells coincides with the positioning of ERGIC marker COPB2 (Fig. 5P).

**Figure 5.**
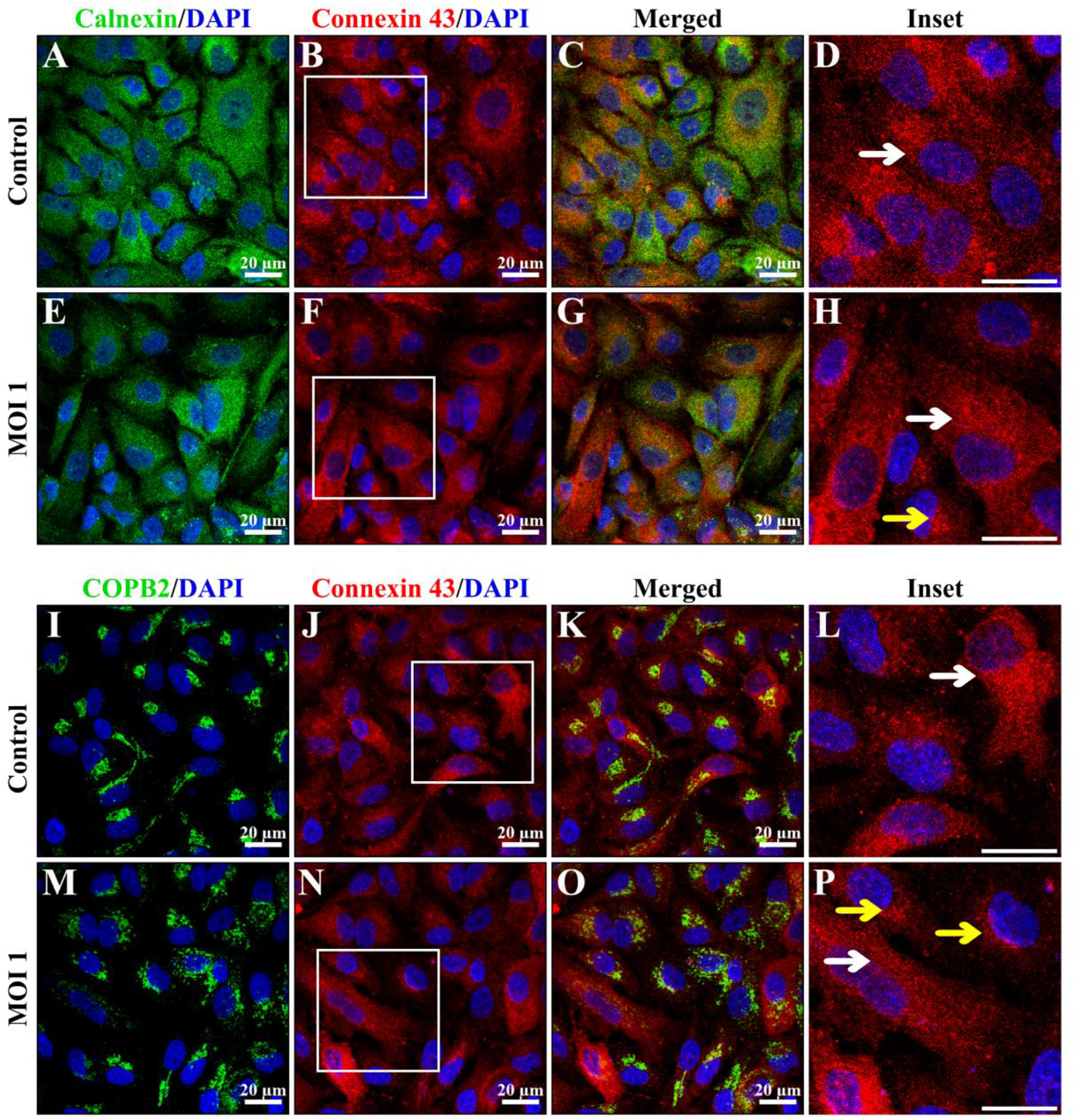
Cx43 localizes in the ERGIC region post OC43 infection in A549 cells. To determine the internal localization of Cx43 post-infection in subcellular compartments, control and infected cells were immunostained with anti-calnexin (A-H) and anti-COPB2 (I-P). The control cells expressing Cx43 staining are marked with white arrows, and the infected cells with perinuclear Cx43 are marked with yellow arrows. Upon visual observation, it is inferred that, upon OC43 infection, Cx43 trafficking is impaired and localizes in the ERGIC compartment of the cells. The insets are digitally zoomed. The scale bar for all the images is 20 microns.

### Functional gap junction and hemichannels are impaired in A549 cells post-HCoV-OC43 infection

Cx43 can function as a gap junction channel and also as a hemichannel. As previously determined upon HCoV-OC43 infection in A549 cells, Cx43 is localized in the ERGIC compartment. Due to the depletion of surface Cx43, gap junction and hemichannel communication gets impaired. Functional gap junction intercellular communication (GJIC) is determined with the help of Lucifer Yellow (LY) dye transfer assay. GJIC allows the passage of molecules less than 1 kDa in weight [23]. LY is a small dye with a molecular weight of around 450 Daltons[24]. LY dye can be transferred from one adjacent cell to another with the help of functional GJIC. We investigated the status of GJIC in control and OC43-infected A549 cells using LY dye transfer assay. Observation in control cells showed that the LY dye was taken up more than OC43 infected cells (Fig. 6A-F). The distance traveled by LY dye from scratch was measured, and the absolute values in microns were plotted (Fig. 6G). From the graph observed it is inferred that the GJIC is impaired in the case of HCoV-OC43 infection. Ethidium Bromide (EtBr) dye uptake assay was performed to identify the presence of functional hemichannels in control and infected A549 cells. The cells were incubated with EtBr in Locke’s solution. The control cells exhibit a brighter EtBr signal (Fig. 6H) than OC43-infected cells (Fig. 6I), depicting a higher level of hemichannel activity (Fig. 6J). In conclusion, the restricted access of Cx43 on the cell surface in HCoV-OC43-infected A549 cells disrupts GJIC and hemichannel activity.

**Figure 6.**
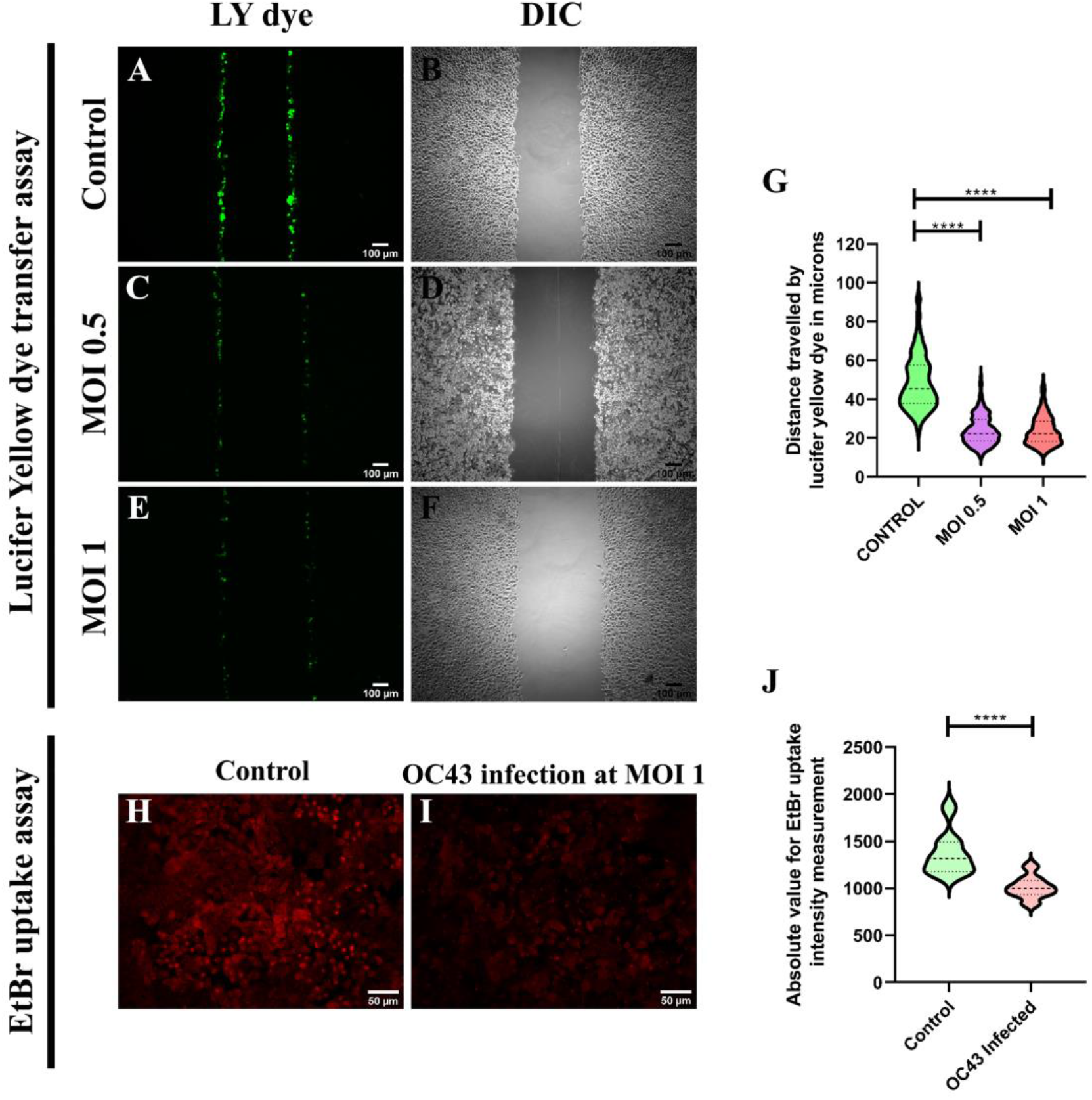
HCoV-OC43 impairs functional GJIC and hemichannel formation. For understanding the status of GJIC functionality, LY dye was scrape loaded in control (A) and MOI 0.5 (C) and 1 (E) OC43 infected cells. It is observed that LY dye uptake and the distance traveled were less in infected A549 cells. Images B, D, and F are DIC images of the respective immunofluorescence fields. The distance traveled by the dye was quantified in Image J, and absolute values were plotted in a violin plot (G). An EtBr uptake assay was performed with control and OC43 infected cells (H and I) to determine hemichannel activity. The absolute value for intensity was measured in Image J and was plotted in a violin plot (J). The scale bar for the LY dye transfer assay images is 100 microns, and all the EtBr uptake assay images are 50 microns. The significant differences are indicated by *P<0.05, **P<0.01, ***P<0.001, ****P<0.0001.

## Discussion

The recent pandemic caused by SARS-CoV-2, a member of the β-coronavirus genus, has left a lasting effect. The lack of research on the pathogenesis of human coronaviruses hindered the urgent need for therapeutics. Therefore, an enhanced understanding of the pathogenesis of human β-coronaviruses is crucial to improving our preparedness for combatting future coronavirus infections. Given the high infectivity of SARS-CoV-2, another human β-coronavirus can be brought under consideration. OC43, also belonging to the β-coronavirus genus, primarily infects the upper respiratory tract and generally manifests a mild clinical condition of common cold [25]. Given the close relationship between SARS-CoV-2 and OC43 within the β-coronavirus family, OC43 can serve as a valuable model for investigating the infection patterns of these human viruses.

This current study aimed to understand the role of human coronaviruses in modulating cell-to-cell communication by focusing on Cx43-mediated gap junction formation. Being expressed ubiquitously in a wide variety of cell lines and playing a crucial role in hemichannel formation facilitating gap junction-mediated communication, Cx43 is an essential part of our study. To the best of our knowledge, this study is the first report showing alteration in Cx43 expression and localization post-HCoV-OC43 infection. These changes directly affect cell-to-cell communication, essential for maintaining cellular homeostasis.

We demonstrated the ability of HCoV-OC43 to infect Vero and A549 cells and its infection spread with respect to day points. With the help of this initial standardization, virus concentrations and incubation time were determined. As mentioned previously, Cx43-mediated cell-to-cell communication is an essential cellular process for maintaining cellular homeostasis. With the help of this study, we aim to understand the modulation of cell-to-cell communications through gap junctions in human β-coronavirus infection. Thus, targeting Cx43 in our research has provided insightful indications for developing therapeutic strategies. In our study, we have selected A549, which is a lung epithelial carcinoma cell line. The primary reason for choosing this cell line for our research is that the virus primarily infects the lungs. Also, the cell line considerably expresses Cx43 both in the cytosol and on the membrane. Therefore, A549 can serve as a good model for studying viral infectivity. HCoV-OC43 infection in A549 cells revealed the downregulation of gap junction protein Cx43 at the protein level. This was demonstrated with the help of Immunoblot and immunofluorescence experiments. This alteration of gap junction protein was accompanied by the increase of other stress markers such as ER stress indicated by HSP70, HSF1, and ERp29 and oxidative stress indicated by DJ-1. Heat shock factor 1 (HSF1) is critical in regulating cellular stress generated by unfolded proteins. Under normal conditions, HSF1 is present as monomers in the cytoplasm. Under stressful situations, HSF1 oligomerizes and retains in the nucleus, where it binds to heat shock elements (HSE) of target genes. One such target gene is Heat shock protein 70 (HSP70). HSP70 is a molecular chaperone expressed under stress that binds to unfolded proteins and stabilizes them. Another ER-resident chaperone that regulates the synthesis and trafficking of several secretory and transmembrane proteins is ERp29. ERp29 is a Thioredoxin-homologous protein with various roles in ER stress and unfolded protein response (UPR). In several viral infections, including SARS-CoV-2, there is oxidative stress in the cells. DJ-1 is known to regulate oxidative stress. It has multiple functions, including antioxidant stress reaction and ROS scavenging [18]. Immunoblot experiments confirmed the upregulation of these stress markers, which further indicates that the internalization of Cx43 may be responsible for increasing the overall stress factors of the cell. Upon infection with OC43, CX43 was observed to be localized in the perinuclear region in infected cells, mostly colocalizing with the ERGIC marker COPB2. The colocalization study was performed using the Immunofluorescence technique, which helped understand the internal localization of Cx43. Triton X-100 solubilization assay confirmed the internal localization of Cx43 in infected cells as infected samples had less Triton X-100 insoluble GJIC plaques. However, we have performed the LY dye transfer assay and EtBr uptake assay to verify if this infectivity negatively regulates functional gap junction formation. The internalization of Cx43 results in alterations of functional gap junction and hemichannel formation, as confirmed by the LY dye transfer assay and EtBr uptake assay, respectively.

Our research has unveiled the pivotal role of viral infections in remodeling intercellular communication. It is clear that intercellular communication is essential for cellular homeostasis. This pathway can prove to be a promising target for developing highly efficacious therapeutics against human coronaviruses by finely regulating the expression of Cx43.

## Acknowledgments

The authors wish to extend their heartfelt appreciation to the Indian Institute of Science Education and Research, Kolkata, for the invaluable support and infrastructure provided. We are grateful to Dr. Michael Koval (Emory University School of Medicine, Atlanta, Georgia, USA) for providing us with the A549 cell line and to Dr. Sourish Ghosh (Indian Institute of Chemical Biology, Kolkata, West Bengal, India) for providing us with Vero cells. We thank Dr. Brenda Hogue (Arizona State University) for providing us with an anti-OC43 antibody. We acknowledge and thank the IISER Kolkata Central Imaging Facility. We thank BEI Resources, NIAID, NIH for the HCoV-OC43 virus. We are grateful to CSIR for providing fellowship to S.K.

## Author Contributions

Conceptualization, J.D.S.; methodology, S.K. and J.D.S. software, S.K.; validation, J.D.S.; formal analysis S.K. and J.D.S.; investigation, S.K., and J.D.S.; resources, J.D.S.; data curation, S.K.; writing—original draft preparation, S.K.; writing—review and editing, S.K. and J.D.S.; visualization, S.K., and J.D.S.; supervision, J.D.S.; project administration, J.D.S.; funding acquisition, J.D.S. All authors have read and agreed to the published version of the manuscript.

## Data Availability Statement

The data are available within this article, its supplementary materials, or from the authors upon request.

